# RBD11, a bioengineered Rab11-binding module for visualizing and analyzing endogenous Rab11

**DOI:** 10.1101/2021.01.25.428185

**Authors:** Futaba Osaki, Takahide Matsui, Shu Hiragi, Yuta Homma, Mitsunori Fukuda

**Author notes:** **Correspondence:** Dr. Takahide Matsui, Dr. Mitsunori Fukuda.

## Abstract

The small GTPase Rab11 plays pivotal roles in diverse physiological phenomena, including the recycling of membrane proteins, cytokinesis, neurite outgrowth, and epithelial morphogenesis. One effective method of analyzing the function of endogenous Rab11 is to overexpress a Rab11-binding domain of one of its effectors, e.g., the C-terminal domain of Rab11-FIP2 (Rab11-FIP2-C), as a dominant-negative construct. However, the drawback of this method is the broader Rab binding specificity of the effector domain, because Rab11-FIP2-C binds to Rabs other than Rab11, e.g., to Rab14 and Rab25. In this study, we bioengineered an artificial Rab11-specific binding domain, named RBD11. Expression of RBD11 visualized endogenous Rab11 without affecting its localization or function, whereas expression of a tandem RBD11, named 2×RBD11, inhibited epithelial morphogenesis and induced a multi-lumen phenotype characteristic of Rab11-deficient cysts. We also developed two tools for temporally and reversibly analyzing Rab11-dependent membrane trafficking: tetracycline-inducible 2×RBD11 and an artificially oligomerized domain (FM)-tagged RBD11.

## INTRODUCTION

Rab GTPases belong to the Ras superfamily of small GTPases and play important roles in membrane trafficking in eukaryotic cells (reviewed in Hutagalung and Novick, 2011; Zhen and Stenmark, 2015; Pfeffer, 2017; Homma et al., 2020). Like other Ras-like GTPases, Rabs cycle between a GTP-bound active state and a GDP-bound inactive state, and the active Rabs promote various membrane trafficking steps, including vesicle budding, tethering, docking, and fusion, through interaction with their specific effectors. Rab cycling is spatiotemporally controlled by two regulatory enzymes: a guanine nucleotide exchange factor and a GTPase-activating protein (reviewed in Ishida et al., 2016; Lamber et al., 2019). Rabs constitute the largest subfamily of the Ras superfamily, and approximately 60 different Rabs have been identified in mammals, as opposed to only 11 Rabs in the unicellular budding yeast (Diekmann et al., 2011; Klöpper et al., 2012). The expansion of Rab isoforms in multicellular eukaryotes, especially in higher eukaryotes, is generally thought to be related to the complexity of their tissues, which consist of highly specialized, differentiated cells that contain unique membrane trafficking pathways. The function of each Rab in mammals has recently been gradually elucidated, but the precise function and localization of most mammalian Rabs remains largely unknown.

Several methods of investigating the function and localization of specific Rabs, such as overexpression of constitutively active or negative (CA/CN) Rab mutants have been developed (reviewed in Fukuda, 2010). One such method uses fluorescently tagged Rab binding (or effector) domains (RBDs) to visualize “endogenous” Rabs and inhibit their functions. However, a drawback of this method is that many of the RBDs bind to several distinct Rabs (Fukuda et al., 2008; Gillingham et al., 2014), and, with few exceptions, their Rab binding specificity has never been thoroughly investigated (Fukuda et al., 2008; 2011; Nottingham et al., 2011; Espinosa et al., 2014; Ohishi et al., 2019). Even the representative effector proteins (e.g., Rabenosyn-5, Rab-interacting lysosomal protein [RILP], and Rab11-FIPs) of the well-characterized, evolutionarily conserved Rabs (e.g., Rab5, Rab7, and Rab11, respectively) bind to several distinct Rabs (Eathiraj et al., 2005; Fukuda et al., 2008; Kelly et al., 2009; Matsui et al., 2012; Schafer et al., 2016). Thus, careful evaluation of their effects on membrane trafficking is necessary when their RBDs are used as dominant-negative constructs, because they can trap several distinct Rabs. Thus, an artificial RBD that can recognize a “single Rab isoform” must be generated by bioengineering techniques to overcome this problem.

In this study, we bioengineered and developed an RBD that is specific for active Rab11 from the C-terminal domain of Rab11-FIP2 and Rab11-FIP4 (Cullis et al., 2002; Hales et al., 2002; Lindsay and McCaffrey, 2002; Wallace et al., 2002) and named it RBD11. We then demonstrated that RBD11 visualized endogenous Rab11 without altering its distribution or single-lumen formation in three-dimensional (3D) cysts formed by Madin-Darby canine kidney (MDCK) cells. We also developed a tandem RBD11, named 2×RBD11, as a dominant-negative construct and showed that expression of 2×RBD11 in MDCK 3D cysts induced a multi-lumen phenotype, the same as occurs in Rab11-deficient cysts (Bryant et al., 2010; Mrozowska and Fukuda, 2016a; Homma et al., 2019). In addition, we developed a tetracycline (Tet)-inducible 2×RBD11 and an artificially oligomerized domain (FM)-tagged RBD11 (Rivera et al., 2000) and demonstrated their usefulness in temporally and reversibly analyzing Rab11-dependent membrane trafficking.

## RESULTS

### Broad and distinct Rab binding specificity of Rab11-FIP1–Rab11-FIP5

To develop a specific RBD that binds to active Rab11 alone, we turned our attention to well-known Rab11 effectors, the Rab11-FIP proteins (reviewed in Horgan and McCaffrey, 2009). Five different Rab11-FIPs (Rab11-FIP1–Rab11-FIP5) have been identified in humans and mice (Figure 1A) and their binding properties of several Rabs such as Rab11 and Rab25 have been well characterized (Prekeris et al., 2000; 2001; Hales et al., 2001; Lindsay et al., 2002; Wallace et al., 2002; Lall et al., 2013; 2015). However, their Rab binding specificity to all mammalian Rabs had never been thoroughly investigated. To comprehensively identify their Rab binding specificity, we cloned cDNAs encoding the C-terminal domain (i.e., Rab-binding domain; RBD) of mouse Rab11-FIP1–Rab11-FIP5 (brackets in Figure 1A) and performed yeast-two hybrid assays using 62 different constitutively negative (CN) Rab mutants (e.g., Rab11A(N124I)) or constitutively active (CA) Rab mutants (e.g., Rab11A(Q70L)) as bait (Figure 1C). Consistent with the results of previous studies, all of the Rab11-FIPs bound to the CA form of Rab11A and Rab11B (red boxes in Figure 1C), but each of them exhibited slightly different Rab binding specificity. In addition, several previously unknown interactions of Rab11-FIPs with Rabs (i.e., Rab20 and Rab42) were also observed, e.g., Rab11-FIP2 bound to the CN form of Rab11A/B and Rab20 (white boxes in Figure 1C) and to the CA form of Rab11A/B, Rab14, and Rab25, whereas Rab11-FIP4 bound to the CA form of Rab11A/B and Rab42. Based on their domain organizations and the sequence similarity of their RBDs (Figure 1A and 1B), Rab11-FIPs have been classified into two groups, class I (Rab11-FIP1, Rab11-FIP2, and Rab11-FIP5) and class II (Rab11-FIP3 and Rab11-FIP4). The class I Rab11-FIPs are characterized by binding to the CA form of Rab14 (yellow boxes in Figure 1C) (Fukuda et al., 2008; Lall et al., 2015), whereas the class II Rab11-FIPs are characterized by binding to the CA form of Rab42 (blue boxes in Figure 1C). We especially noted that all of the Rab11-FIPs except Rab11-FIP4-C bound to the CN form of Rab11A and Rab11B (orange boxes in Figure 1C) and to the GDP-bound form of Rab11 (Junutula et al., 2004). Rab11-FIP2 also bound to Rab11A/B(S25N), another CN form of Rab11A/B (Hales et al., 2001; Lindsay and McCaffrey, 2002), whereas Rab11-FIP4 did not (Figure S1A and S1B). In view of its narrow Rab binding specificity and strict GTP-dependency, we decided to use Rab11-FIP4-C as the main backbone to develop an active Rab11-specific binding module.

**Figure 1.**
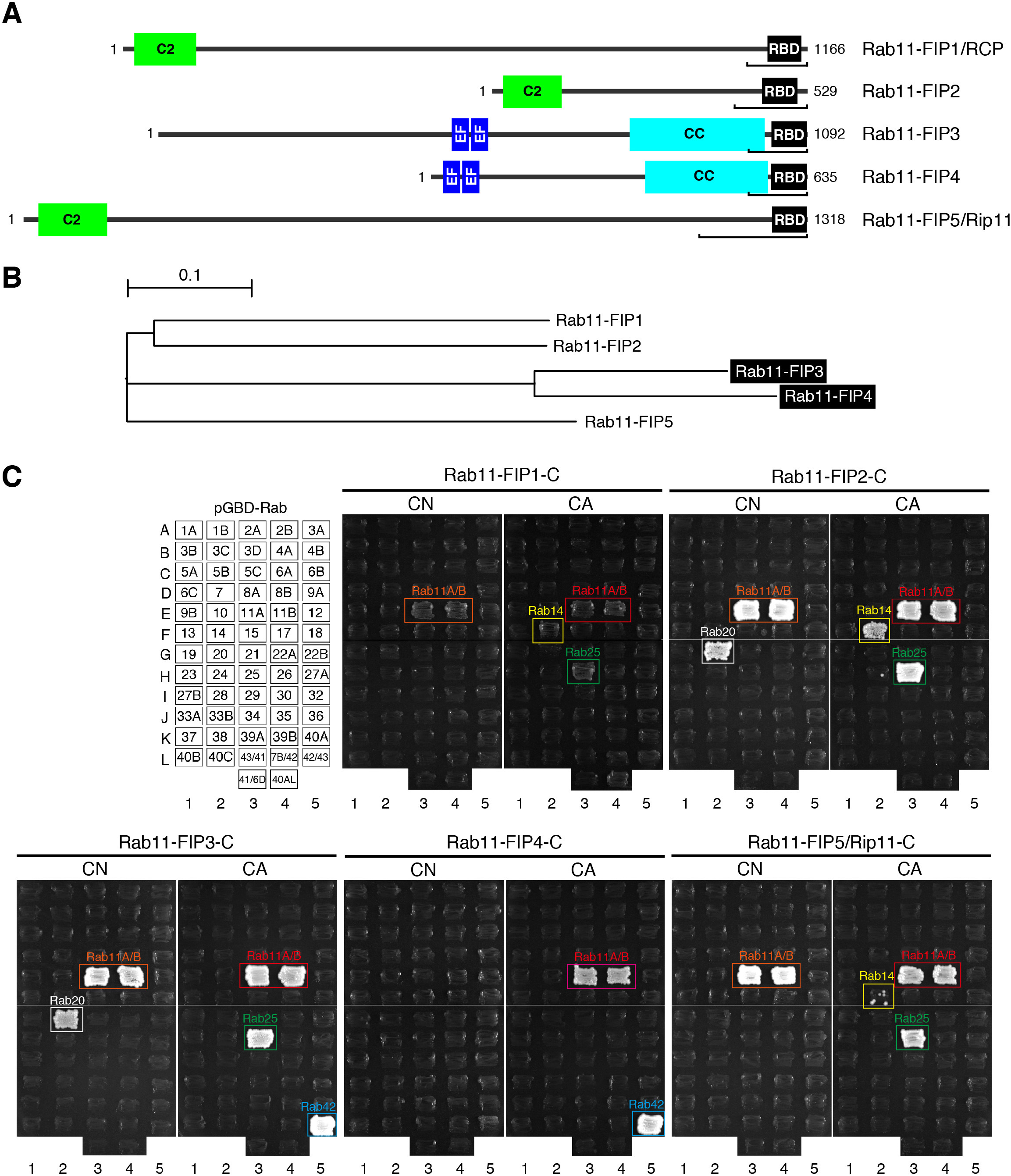
Distinct Rab binding specificity of mouse Rab11-FIP proteins as revealed by yeast two-hybrid assays. (A) Schematic representation of mouse Rab11-FIPs, which share a C-terminal Rab-binding domain (RBD). Class I Rab11-FIPs (i.e., FIP1, FIP2, and FIP5) contain an N-terminal C2 domain, whereas class II Rab11-FIPs (i.e., FIP3 and FIP4) contain EF hand domains and a coiled-coil (CC) domain just before the RBD. The brackets indicate the regions used to perform the yeast two-hybrid assays in (C). (B) A phylogenetic tree of the RBD of mouse Rab11-FIPs. The class II Rab11-FIPs are shown on a black background. (C) Rab binding specificity of the RBD of mouse Rab11-FIPs as determined by yeast two-hybrid assays. The C-terminal region of Rab11-FIPs was subcloned into the pGAD-C1 vector, and it was then transformed into yeast cells expressing pGBD-C1 vector carrying a CN or CA form of each Rab (positions are indicated in the upper left panel). Interactions were detected by the growth of the yeast cells, and positive patches are boxed: Rab11(CN) in orange, Rab20(CN) in white, Rab11(CA) in red, Rab14(CA) in yellow, Rab25(CA) in green, and Rab42(CA) in blue.

### Development of an engineered Rab-binding domain specific for Rab11 (RBD11)

Since the class II Rab11-FIPs bound to Rab42 in addition to Rab11 (Figure 1C), we next investigated whether Rab42 binds to the Rab11-binding site of Rab11-FIP4-C (or Rab11-FIP3-C). To do so, we deleted one third of the C-terminal portion of Rab11-FIP4-C (or Rab11-FIP3-C) (Figure 2A and see also Figure 3A), which is known to be essential for Rab11 binding (Eathiraj et al., 2006; Jagoe *et al*., 2006; Shiba et al., 2006), and tested the Rab11/42 binding ability of Rab11-FIP3-ΔC and Rab11-FIP4-ΔC interacted with neither Rab11A/B nor Rab42, suggesting that Rab11 and Rab42 bind to the same region of Rab11-FIP3 and Rab11-FIP4. To artificially alter the Rab binding specificity of Rab11-FIP4-C, we then prepared two chimeric proteins between Rab11-FIP2-C and Rab11-FIP4-C (black bars and gray bars, respectively, in Figure 2A). Although Rab11-FIP4/2-C2 completely lacked Rab11/42 binding ability, Rab11-FIP4/2-C1 bound to the CA form of Rab11A/B, but not to the CA form of Rab42 (Figure 2C). Fortunately, Rab11-FIP4/2-C1 bound strongly to active Rab11A/B and weakly to Rab25 (sometimes called Rab11C), but it did not bind to inactive Rabs (Figure 2D).

**Figure 2.**
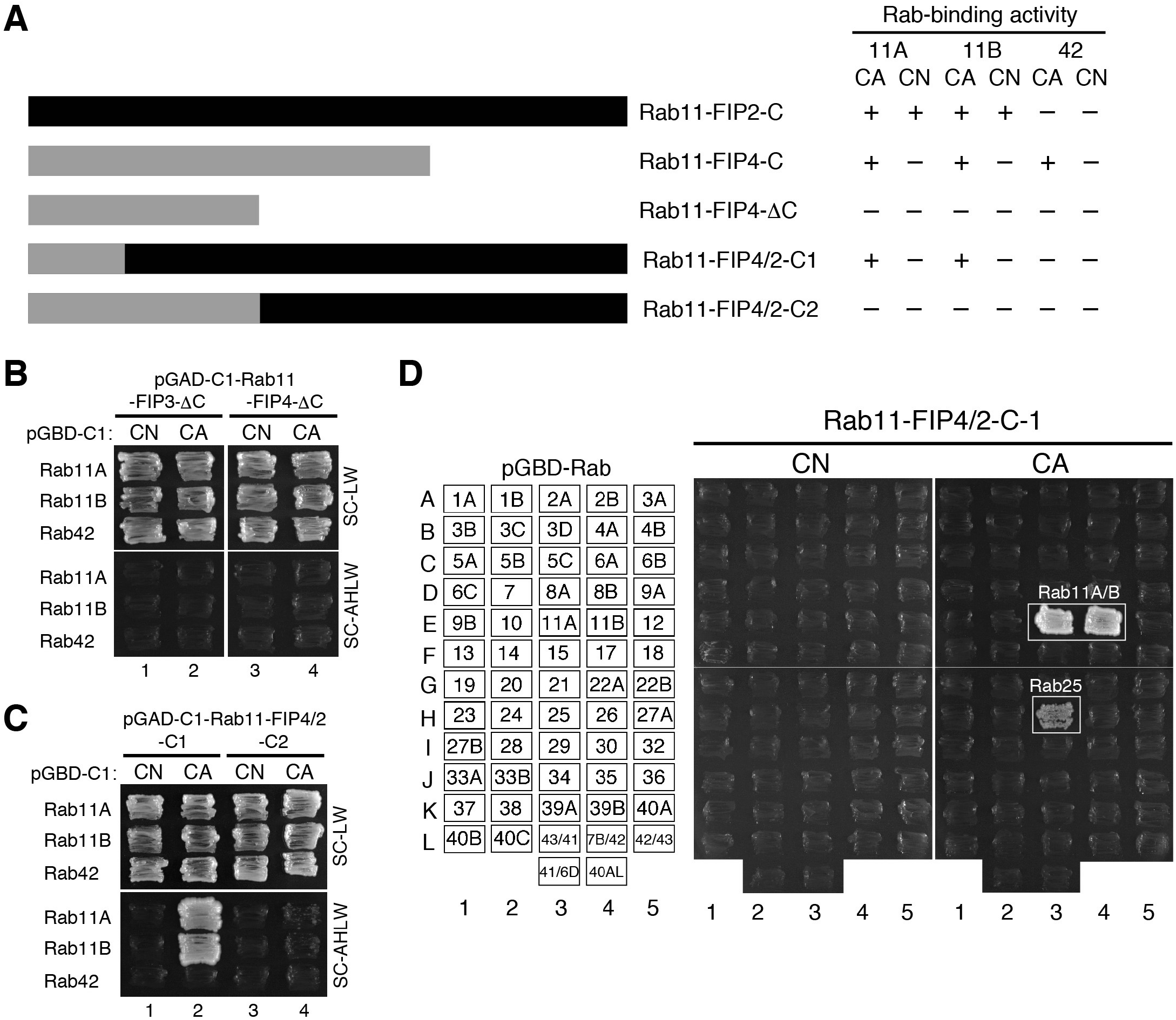
Increased Rab11 binding specificity of a chimeric construct between Rab11-FIP4 and Rab11-FIP2 (Rab11-FIP4/2-C1) (A) Schematic representation of the deletion mutant of Rab11-FIP4-C and chimeric constructs between Rab11-FIP4-C (gray bars) and Rab11-FIP2-C (black bars) used in this study (see also Figure 3A). (B) Rab binding specificity of the C-terminal deletion mutants of Rab11-FIP3 and-FIP4 (Rab11-FIP3-ΔC and FIP4-ΔC) as determined by yeast two-hybrid assays. Yeast cells were grown on SC-LW (growth medium) and SC-AHLW (selection medium). (C) Rab binding specificity of the chimeric proteins, Rab11-FIP4/2-C1 and-C2 as determined by yeast two-hybrid assays. (D) Rab binding specificity of Rab11-FIP4/2-C1 as determined by yeast two-hybrid assays. See also the legend to Figure 1C.

**Figure 3.**
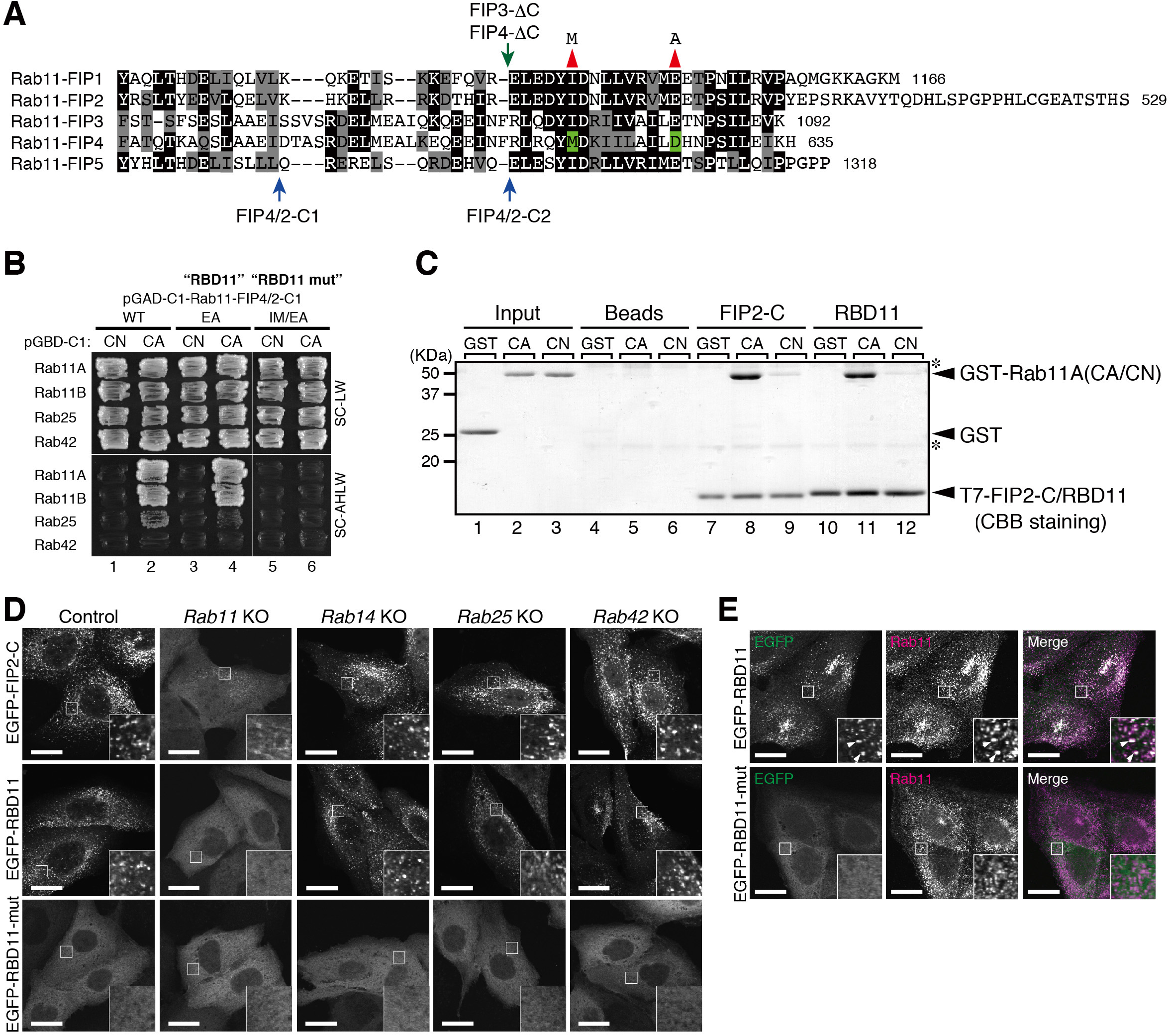
Development of a novel Rab11-binding domain (RBD11) (A) Sequence alignment of the C-terminus of mouse Rab11-FIPs. Identical and similar amino acids in more than half of the sequences are indicated by displaying them against a black background and gray background, respectively. The green arrow points to the deletion sites in Rab11-FIP3-ΔC and FIP4-ΔC in Figure 2A and 2B. The blue arrows point to the chimeric sites of Rab11-FIP4/2-C1 and-C2 in Figure 2A and 2C. The red arrowheads indicate the position of the amino acid substitution in RBD11 (i.e., Ile-to-Met substitution) and RBD11-mut (i.e., Ile-to-Met and Glu-to-Ala substitutions). (B) Rab binding specificity of Rab11-FIP4/2-C1 and its point mutants (EA and EA/IM) as determined by yeast two-hybrid assays. Yeast cells were grown on SC-LW (growth medium) and SC-AHLW (selection medium). (C) GTP-dependent interaction between T7-tagged Rab11-FIP2-C or RBD11 and GST-Rab11 (CA or CN) as revealed by direct binding assays. The asterisks indicate nonspecific bands that originated from the anti-T7-tag antibody (i.e., light and heavy chains of IgG). (D) Subcellular localization of EGFP-tagged Rab11-FIP2-C, RBD11, and RBD11-mut in *Rab11*-KO, *Rab14*-KO, *Rab25*-KO, and *Rab42*-KO MDCK cells. Note that RBD11, not Rab11-FIP2-C, yielded a complete diffuse cytosolic pattern only in *Rab11* KO cells. (E) Subcellular localization of EGFP-tagged RBD11 and RBD11-mut in wild-type MDCK cells. Note that RBD11, not RBD11-mut, colocalized well with endogenous Rab11.

To further manipulate the Rab25 binding ability of Rab11-FIP4/2-C1, we performed site-directed mutagenesis, especially focusing on amino acids that are conserved only in Rab11-FIP4, because Rab11-FIP4-C alone did not bind to Rab25 in our two-hybrid assays (green boxes in Figure 1C). Sequence comparisons of the RBDs of Rab11-FIPs enabled us to identify Met-616 (corresponding to Ile-481 of Rab11-FIP2) and Asp-625 (corresponding to Glu-490 of Rab11-FIP2) as unique residues in Rab11-FIP4 (green background in Figure 3A). When the Glu residue of Rab11-FIP4/2-C1 (red arrowhead on the right in Figure 3A) was mutated to Ala, the resulting Rab11-FIP4/2-C1-EA mutant hardly interacted with Rab25(CA) (lane 4 in Figure 3B). When the Ile residue of Rab11-FIP4/2-C1 that corresponds to Ile-480 of Rab11-FIP2, which is essential for Rab11 interaction (Jagoe et al., 2006), was further mutated to Met (red arrowhead on the left in Figure 3A), the resulting Rab11-FIP4/2-C1-IM/EA mutant completely eliminated its Rab11/25(CA) binding ability (lane 6 in Figure 3B). We therefore decided to use Rab11-FIP4/2-C1-EA as “RBD11 (RBD specific for active Rab11)” and Rab11-FIP4/2-C1-IM/EA (referred to as RBD11-mut hereafter) as an ideal negative control for RBD11.

To confirm that RBD11 directly binds to Rab11 in a GTP-dependent manner, we performed direct binding assays using purified components (T7-tagged RBD11 and GST-Rab11A(CA/CN)) (Figure 3C). As anticipated, T7-RBD11 strongly bound to GST-Rab11A(CA), and hardly bound to GST-Rab11A(CN) at all (lanes 11 and 12 in Figure 3C). Although T7-Rab11-FIP2-C also preferentially bound to GST-Rab11(CA) based on the results of the precipitation assays, it always bound to GST-Rab11(CN) more strongly than RBD11 did (lanes 9 and 12 in Figure 3C), consistent with the results of the yeast two-hybrid assays described above. Since in addition to binding to Rab11, Rab11-FIP4 has been reported to bind to Arf6, another type of Ras-like small GTPase (Fielding et al., 2005; Shiba et al., 2006), we also confirmed by yeast two-hybrid assays that RBD11 does not trap active Arf6 (Figure S1C).

Next, we investigated whether RBD11 also selectively binds to Rab11 in cultured mammalian cells by using a recently established *Rab* KO collection (Homma et al., 2019), because we think that the results obtained above in yeast cells may not be simply applied to the Rab binding specificity of RBD11 in mammalian cells. When EGFP-tagged Rab11-FIP2-C, RBD11, and RBD11-mut were each stably expressed in wild-type (control), *Rab11A/B*-KO (*Rab11*-KO), *Rab14*-KO, *Rab25*-KO, and *Rab42*-KO MDCK cells, both Rab11-FIP2-C and RBD11 showed a punctate distribution in the control cells, whereas RBD11-mut exhibited a cytosolic distribution (far left column in Figure 3D). It should be noted that *Rab11*-KO cells were the only *Rab*-KO cells in which RBD11 yielded a complete cytosolic distribution (middle row in Figure 3D). By contrast, a punctate distribution of Rab11-FIP2-C was still observed even in *Rab11*-KO cells, although its punctate signals were clearly decreased (top row in Figure 3D), suggesting that Rab11-FIP2-C traps Rabs other than Rab11A/B in cultured mammalian cells. We then immunostained for endogenous Rab11 with a specific antibody and confirmed that RBD11, not RBD11-mut, colocalized with endogenous Rab11 (Figure 3E). Consistent with the recycling endosomal localization of Rab11, RBD11 did not colocalize with any other organelle markers, including GM130 (Golgi marker), EEA1 (early endosome marker), LBPA (late endosome marker), and LAMP2 (lysosome marker) (Figure S2A).

### Development of a tandem RBD11 (2×RBD11) capable of inhibiting Rab11-dependent membrane trafficking during MDCK 3D cyst formation

To investigate the effect of RBD11 on Rab11-dependent membrane trafficking events, we focused on single-lumen formation by 3D cysts formed by MDCK cells, because a multi-lumen phenotype was clearly observed in *Rab11*-KO cysts and *Rab11*-knockdown (KD) cysts (far left image in Figure 4B) (Bryant et al., 2010; Homma et al., 2019). We prepared 3D MDCK cysts stably expressing EGFP-tagged RBD11, RBD11-mut, or Rab11-FIP2-C and visualized their luminal domain (i.e., apical domain) by staining with anti-ezrin antibody. Since both Rab11-FIP2-C and RBD11 recognized endogenous Rab11 (Figure 3D and 3E), we initially expected that their expression should induce a typical multi-lumen phenotype by trapping endogenous Rab11 (i.e., by serving as a “Rab11 trapper”). Actually, many cysts stably expressing EGFP-Rab11-FIP2-C contained multiple small lumens (upper right image in Figure 4B), but contrary to our expectations, stable expression of EGFP-RBD11 (or EGFP-RBD11-mut) failed to impair single lumenogenesis (lower panels in Figure 4B).

**Figure 4.**
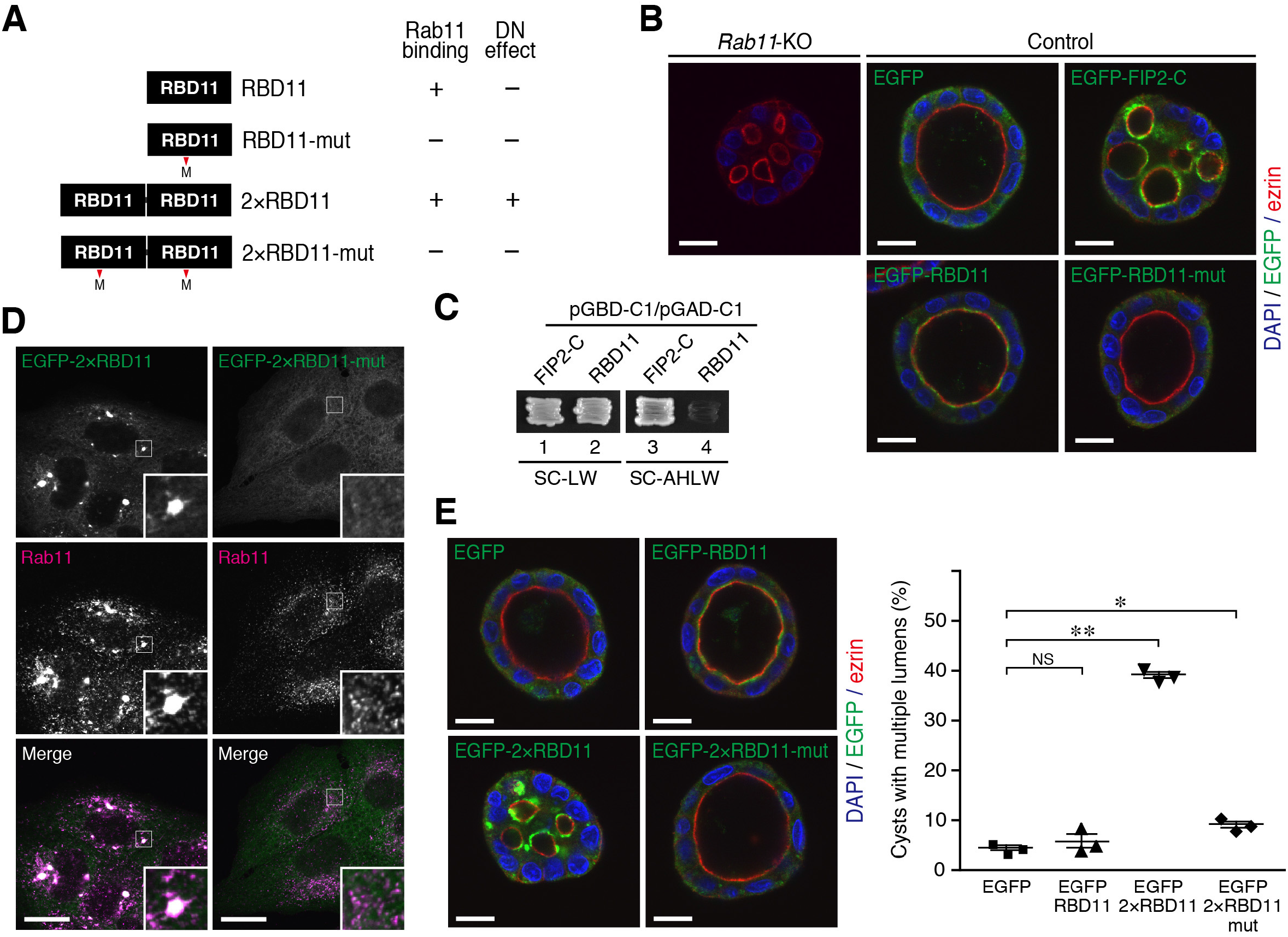
2×RBD11 serves as an effective Rab11 trapper. (A) Schematic representation of RBD11, 2×RBD11, and their mutants lacking Rab11 binding ability. DN, dominant-negative. (B) Typical images of Rab11-KO cysts and wild-type (control) MDCK cysts stably expressing EGFP-tagged Rab11-FIP2-C, RBD11, or RBD11-mut. Each cyst was cultured in collagen gel for 7 days. The cysts were then fixed with 10% TCA and stained with anti-EGFP (green) and anti-ezrin (red) antibodies. Scale bars, 20 μm. (C) Self-oligomerization activity of Rab11-FIP2-C and RBD11 as determined by yeast two-hybrid assays. Yeast cells expressing the vectors indicated were grown on SC-LW (growth medium) and SC-AHLW (selection medium). (D) Subcellular localization of 2×RBD11 and 2×RBD11-mut in wild-type MDCK cells. Cells stably expressing EGFP-tagged 2×RBD11 or 2×RBD11-mut (green) were fixed with 4% PFA and immunostained with anti-Rab11 antibody (red). (E) Typical images of wild-type MDCK cysts stably expressing EGFP, EGFP-tagged RBD11, 2×RBD11, or 2×RBD11-mut. Cysts were cultured in collagen gel for 7 days, fixed with 10% TCA, and then stained with anti-EGFP (green) and anti-ezrin (red) antibodies. Scale bars, 20 μm. The graph on the right shows the percentage of cysts containing multiple lumens. Data are means and SEM from three independent experiments (200 cysts per experiment). **, *p* <0.001; *, *p* <0.05; N.S., not significant (one-way ANOVA and Tukey’s test).

To determine why RBD11 failed to affect single-lumen formation, we focused on another biochemical property of Rab11-FIP2-C, i.e., its dimerization activity (Lindsay and McCaffrey, 2002; Junutula et al., 2004; Jagoe et al., 2006; Wei et al., 2006), because a dimerized protein would trap its ligand more efficiently by increasing its local concentration. Consistent with the results of previous studies, the results of yeast two-hybrid assays showed that Rab11-FIP2-C formed homodimer, but that RBD11 did not exhibit homodimerization activity (compare lanes 3 and 4 in Figure 4C). We therefore attempted to confer a dominant-negative function on RBD11 by arranging RBD11 in tandem (named 2×RBD11) (Figure 4A). When EGFP-tagged 2×RBD11 was stably expressed in MDCK cells, large Rab11-positive puncta, which also colocalized with EGFP-2×RBD11 (insets in the left column of Figure 4D), were observed in the perinuclear region. Moreover, these puncta did not colocalize with other organelles, including the Golgi apparatus, early endosomes, late endosomes, and lysosomes, where Rab11 was not present (Figure S2B). By contrast, no such large puncta were observed in EGFP-2×RBD11-mut-expressing cells (right column of Figure 4D). In contrast to the original EGFP-RBD11 and EGFP-2×RBD11-mut, EGFP-2×RBD11 was found to significantly inhibit single-lumen formation by 3D cysts (Figure 4E).

### Temporal inhibition of Rab11 by Tet-inducible 2×RBD11 and artificially oligomerized RBD11 (FM-RBD11)

Finally, we attempted to create two additional RBD11-based tools capable of temporally inhibiting the function of endogenous Rab11. First, we established MDCK cell lines, in which expression of 2×RBD11 (or 2×RBD11-mut as a control) was specifically induced by Tet (Figure 5A, upper two constructs) and confirmed its doxycycline (Dox)-inducible expression (Figure 5B). We then used the Tet-inducible system to investigate the possible involvement of Rab11 in single-lumen formation by 3D cysts and to determine the stage at which Rab11 functions during lumenogenesis. As shown in Figure 5C and 5D, Tet-induced expression of 2×RBD11 in the first half of cyst growth (i–iv) resulted in a significant increase in the number of cysts containing multiple lumens, whereas its expression in the latter half did not (v). By contrast, Tet-induced expression of 2×RBD11-mut had no effect on lumenogenesis under any conditions (Figure 5C), indicating that the multi-lumen phenotype induced by 2×RBD11 in the first half of cyst growth is attributable to inhibition of endogenous Rab11.

**Figure 5.**
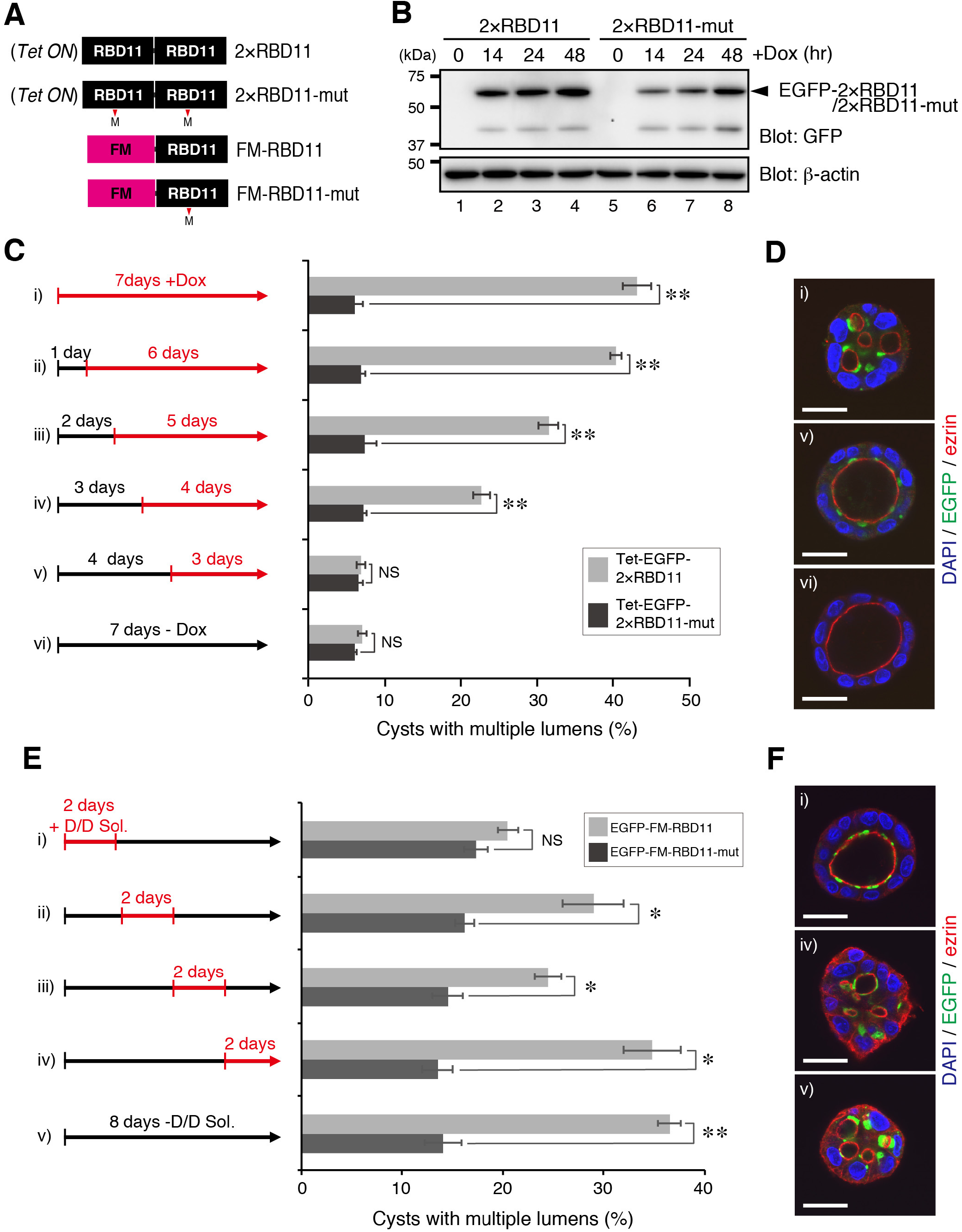
Inducible expression of 2×RBD11 or FM-RBD11 had a DN effect on MDCK 3D cyst formation. (A) Schematic representation of tetracycline-inducible (Tet-ON) 2×RBD11, FM-tagged RBD11, and their mutants lacking Rab11 binding ability. FM, a FKBP12-derived artificial oligomerization domain. (B) Doxycycline (Dox)-induced expression of EGFP-2×RBD11 was detected by immunoblotting. Lysates from MDCK cells stably expressing Tet-ON-2×RBD11 that had been treated with 2 μg/mL Dox for the times indicated were immunoblotted with anti-GFP and anti-β-actin antibodies. (C) EGFP-tagged 2×RBD11 or 2×RBD11-mut under the control of Tet-ON promoters was expressed in MDCK cysts, and the percentages of cysts containing multiple lumens were calculated. Cells were cultured in collagen gel for 7 days in the presence of 2 μg/mL Dox (red lines) for the times indicated, fixed with 10% TCA, and then stained with anti-GFP (green) and anti-ezrin (red) antibodies. Data are means and SEM from three independent experiments (200 cysts per experiment). **, *p* <0.001; *, *p* <0.05; N.S., not significant (two-sided Student’s unpaired *t*-test). (D) Typical images of cysts shown in (C)-i, −v, and vi. Scale bars, 20 μm. (E) EGFP-tagged FM-RBD11 or FM-RBD11-mut was expressed in MDCK cysts, and the percentages of cysts containing multiple lumens were calculated. Cells were cultured in collagen gel for 8 days in the presence of 250 nM D/D solubilizer (red lines) for the times indicated. Data are means and SEM from three independent experiments (200 cysts per experiment). *, *p* <0.05; **, *p* <0.001; N.S., not significant (two-sided Student’s unpaired *t*-test). (F) Typical images of cysts shown in (E)-i, iv, and v. Scale bars, 20 μm.

Because 2×RBD11, not RBD11 alone, inhibited single-lumen formation by 3D cysts (Figure 4E), we assumed that artificial regulation of RBD11 dimerization (i.e., a rapid transition between monomer and dimer) in living cells would allow us to temporally and reversibly inhibit the function of endogenous Rab11. To test our assumption, we focused on a drug-regulated homodimerization domain (i.e., FM domain) (Rivera et al., 2000) and prepared FM-tagged RBD11 and RBD11-mut (Figure 5A, lower two constructs). In the absence of D/D solubilizer, FM-RBD11 forms a dimer, which presumably inhibits the function of endogenous Rab11, the same as 2×RBD11 does. By contrast, in the presence of D/D solubilizer, FM-RBD11 is a monomer, which is unlikely to have any effect on the function of Rab11. We therefore treated MDCK cells stably expressing FM-RBD11 (or FM-RBD11-mut) with D/D solubilizer for the times indicated in Figure 5E and examined its effect on lumenogenesis by 3D cysts. Consistent with the results obtained with the Tet-inducible 2×RBD11 described above, treatment of FM-RBD11-expressing cells with D/D solubilizer only during the first two days of cyst growth completely reversed the inhibitory effect of FM-RBD11, and the cells showed normal single lumenogenesis (Figure 5E(i) and 5F(i)). By contrast, when cells were exposed to D/D solubilizer after two days of cyst growth, a significantly increased number of cysts containing multiple lumens was observed even in the presence of D/D solubilizer (Figure 5E(ii)-(iv) and 5F(iv)). Again, the Rab11-binding-deficient FM-RBD11-mut had no effect on lumenogenesis under any conditions (Figure 5E). All of these findings were highly consistent with the results of previous studies showing that apical membrane proteins are transcytosed to the newly formed apical domain through Rab11-positive endosomes in the early stage of cyst formation (Marc et al., 2009) and that Rab11 mediates the formation of a single apical membrane initiation site (Bryant et al., 2010; Mrozowska and Fukuda, 2016a). Since these Rab11-dependent events occur 16–36 hr after the start of cyst growth (Mrozowska and Fukuda, 2016b), it is reasonable to expect that inhibition of endogenous Rab11 with 2×RBD11 or FM-RBD11 during the initial 48-hr period of cyst growth is the most effective means of inducing a multi-lumen phenotype.

## DISCUSSION

In the present study, we performed mutational and chimeric analyses of Rab11-FIP2 and Rab11-FIP4 and succeeded in developing an artificial protein module named RBD11 that specifically binds to the active Rab11 isoforms, i.e., Rab11A and Rab11B, both *in vitro* and in cultured cells. We think that the RBD11 we developed has several advantages over the RBD of Rab11-FIP2. The first advantage is its exclusive Rab11 binding specificity and specific recognition of the GTP-bound form of Rab11 (Figure 3). The second advantage is that a strict negative control for RBD11, named RBD11-mut, a point mutant of RBD11 that completely lacks Rab11 binding ability, can be used to determine the subcellular localization (Figure 3D and 3E) and inhibit the function of endogenous Rab11 (Figure 4E). The third advantage is that since, in contrast to the original Rab11-FIP2-C construct (Figure 4B), stable expression of RBD11 itself in MDCK cells had no effect on the function of Rab11, it can be used as a tool to visualize endogenous, active Rab11 *in vivo* without altering its localization or inhibiting its function. It should be noted, however, that tandem RBD11 (2×RBD11) and artificially dimerized RBD11 (FM-RBD11) significantly inhibited single-lumen formation by 3D MDCK cysts, thereby leading to a multi-lumen phenotype (Figures 4E and 5), which is characteristic of Rab11-deficient cysts (Figures 4B). Thus, Tet-inducible 2×RBD11 and artificially dimerized RBD11 are unique tools that can be used to temporally and reversibly analyze the function of Rab11 at the endogenous protein level.

Several methods for analyzing Rab11-mediated membrane trafficking have become available thus far. The most widely used method is overexpression of a CA or CN form of Rab11. However, a drawback of this method is that CA/CN Rabs have sometimes affected membrane trafficking even though the corresponding Rabs have not been endogenously expressed. By contrast, KD of *Rab11* with a specific siRNA and KO of *Rab11* by genome-editing technologies are powerful methods for analyzing the function of endogenous Rab11. However, because two Rab11 isoforms, Rab11A and Rab11B, are present in mammals and function redundantly, at least in epithelial morphogenesis by MDCK cells (Homma et al., 2019), simultaneous KD or KO of *Rab11A/B* is necessary, and KD/KO efficiency is a limiting factor. Moreover, off-target effects of siRNA or guide RNA should also be considered, and appropriate rescue experiments are generally required. In that sense, the dimeric form of FM-RBD11 and monomeric form of FM-RBD11 (or RBD11 and RBD11-mut) developed in this study are ideal positive and negative controls, respectively, for analyzing the localization and function of endogenous Rab11. A Förster resonance energy transfer (FRET)-based Rab11 sensor, named AS-Rab11, has previously been reported as a means of visualizing Rab11 activation and inactivation *in vivo* (Campa et al., 2018). Although AS-Rab11 enables spatiotemporal visualization of Rab11 activation and inactivation, it is incapable of inhibiting the function of endogenous Rab11. Conversely, RBD11 is unable to visualize Rab11 activation and inactivation of Rab11 (i.e., it only visualizes active Rab11), but FM-RBD11 is capable of temporally inhibiting the function of endogenous Rab11. Similarly, optogenetically oligomerized Rab11 has been reported to inhibit the function of Rab11 (Nguyen et al., 2016). Although this tool is superior to FM-RBD11 in terms of spatial regulation, it does not directly inhibit the function of endogenous Rab11 and instead inhibits the trafficking of recycling endosomes, where ectopically expressed oligomerized Rab11 is present. Based on all of the above findings taken together, we think that the RBD11 tools developed in this study will serve as powerful tools for initial assessments of the function of endogenous Rab11 in many cell types, because they are easily expressed by means of plasmid transfection or retrovirus infection without the need for any special equipment. However, use of different Rab11 tools, including RBD11, in combination will certainly be necessary to fully understand the spatiotemporal regulation of Rab11-mediated membrane trafficking.

In conclusion, we have bioengineered a Rab11-specific binding module, named RBD11, and further developed FM-RBD11, which is capable of visualizing endogenous Rab11 in the monomer state and inhibiting the function of endogenous Rab11 in the dimer state. Because Rab11 is highly conserved in vertebrates, our RBD11 tools would be applied to various mammalian cell lines, and even to animal models. Our strategy may also apply to other Rabs by using their specific effector domains with FM-tag, which will contribute to the elucidation of the molecular mechanisms of Rab-mediated membrane trafficking in the future.

## TRANSPARENT METHODS

### Materials

The following antibodies were obtained commercially: anti-Rab11 rabbit polyclonal antibody (Invitrogen, Carlsbad, CA; #71-5300), which recognizes both Rab11A and Rab11B, anti-GM130 mouse monoclonal antibody (BD Biosciences, San Jose, CA; #610823), anti-EEA1 mouse monoclonal antibody (BD Biosciences; #610456), anti-LBPA mouse monoclonal antibody (Applied Biological Materials, Richmond, BC, Canada; #G043), anti-LAMP2 mouse monoclonal antibody (Thermo Fisher Scientific, Waltham, MA; #MA-28269), anti-TfR mouse monoclonal antibody (Invitrogen; #13-6800), anti-β-actin mouse monoclonal antibody (Applied Biological Materials; #G043), anti-ezrin mouse monoclonal antibody (Abcam, Cambridge, UK; #ab4069), horseradish peroxidase (HRP)-conjugated anti-GFP polyclonal antibody (MBL, Nagoya, Japan; #598-7), HRP-conjugated anti-mouse IgG goat polyclonal antibody (SouthernBiotech, Birmingham, AL; #1031-05), Alexa Fluor 555^+^-conjugated anti-mouse IgG goat polyclonal antibody (Thermo Fisher Scientific; #A32727), and Alexa Fluor 555^+^-conjugated anti-rabbit IgG goat polyclonal antibody (Thermo Fisher Scientific; #A32732). Other reagents used in this study were also obtained commercially: doxycycline (FUJIFILM Wako Pure Chemical Corporation, Osaka, Japan) and D/D solubilizer (Takara Bio, Shiga, Japan).

### cDNA cloning and plasmid constructions

cDNAs encoding the C-terminal 102 amino acids (AA) of mouse Rab11-FIP1/RCP, the C-terminal 124 AA of mouse Rab11-FIP2, the C-terminal 100 AA of Rab11-FIP3, the C-terminal 99 AA of mouse Rab11-FIP4, and the C-terminal 184 AA of mouse Rab11-FIP5/Rip11 were amplified from the Marathon-Ready adult mouse brain and testis cDNAs (Clontech/Takara Bio) by performing PCR using the standard molecular biology techniques. After verifying their sequences, they were subcloned into the pGAD-C1 vector (James et al., 1996) to perform yeast two-hybrid assays. Deletion mutants of Rab11-FIP3 and Rab11-FIP4 (ΔC), chimeric mutants between Rab11-FIP2 and Rab11-FIP4 (FIP4/2-C1 and FIP4/2-C2), FIP4/2-C2 point mutants (EA [= RBD11] and IM/EA [= RBD11 mut]; see Figure 3A for details) were also prepared by the standard molecular biology techniques, including the PCR sewing technique. The Rab11-FIP2-C, RBD11, and RBD-mut cDNA fragments were subcloned into the pEGFP-C1 vector (Clontech/Takara Bio), pEGFP-C1-FM vector (Hirano et al., 2016), pMRX-IRES-puro-EGFP vector (a kind gift from Dr. Shoji Yamaoka, Tokyo Medical and Dental University, Tokyo, Japan) (Saitoh et al., 2003), pRetroX-TetOne-Puro vector (Clontech/Takara Bio), and/or pEF-T7 tag vector (Fukuda et al., 1999).

cDNAs encoding the following CN mutants of mouse or human Rabs were also produced by the standard molecular biology techniques: Rab1A(N124I), Rab1B(N121I), Rab2A(N119I), Rab2B(N119I), Rab3A(N135I), Rab3B(N135I), Rab3C(N143I), Rab3D(N135I), Rab4A(N126I), Rab4B(N121I), Rab5A(N133I), Rab5B(N147I), Rab5C(N134I), Rab6A(N126I), Rab6B(N126I), Rab6C(N126I), Rab41/6D(N144I), Rab7(N125I), Rab7B/42(N124I), Rab8A(N121I), Rab8B(N121I), Rab9A(N124I), Rab9B(N124I), Rab10(N122I), Rab11A(N124I), Rab11B(N124I), Rab12(N154I), Rab13(N121I), Rab14(N124I), Rab15(N121I), Rab17(N132I), Rab18(N122I), Rab19(N130I), Rab20(N113I), Rab21(N130I), Rab22A(N118I), Rab22B(N118I), Rab23(N121I), Rab24(T120I), Rab25(N125I), Rab26(N181I), Rab27A(N133I), Rab27B(N133I), Rab28(N129I), Rab29(N125I), Rab30(N122I), Rab32(N141I), Rab33A(N151I), Rab33B(N148I), Rab34(S166I), Rab35(N120I), Rab36(T171I), Rab37(N143I), Rab38(N127I), Rab39A(H127I), Rab39B(H123I), Rab40A(N126I), Rab40AL(N126I), Rab40B(N126I), Rab40C(N126I), Rab43/41(N129I), and Rab42/43(H127I). The nomenclature of the Rabs in this study is in accordance with the National Center for Biotechnology Information (NCBI) database, and the names of several Rabs in the report by Itoh *et al*. (2006) are different (indicated by slash in Figure 1C). The Rab CN mutants lacking a 3’ region that encodes Cys residue(s) for geranylgeranylation were subcloned into the pGBD-C1 vector (named pGBD-C1-Rabs(CN)ΔCys; James et al., 1996). Mouse Arf6-Q67L (CA form) and Arf6-T27N (CN form) cDNA fragments (Kobayashi and Fukuda, 2012) were also subcloned into the pGBD-C1 vector. pGBD-C1-Rabs(CN)ΔCys vectors were prepared as described previously (Fukuda et al., 2008). pGBD-C1-Rab11A(S25N)ΔCys, −Rab11B(S25N)ΔCys, −Rab14(S25N)ΔCys, −Rab25(T26N)ΔCys, and −Rab42(T23N)ΔCys were also prepared as described previously (Tamura et al., 2009). Mouse Rab11A(Q70L) (= CA; Itoh et al., 2006) and Rab11A(N124I) (= CN) cDNA fragments were subcloned into the pGEX-4T-3 vector (GE Healthcare, Buckinghamshire, UK). The sequences of the oligonucleotides used for plasmid constructions in this study are available from the corresponding authors on request.

### Yeast two-hybrid assays

The yeast strain (PJ69-4A), medium, culture conditions, and transformation protocol used were as described previously (James et al., 1996). The yeast two-hybrid assays were performed using pGBD-C1-Rabs(CA/CN)ΔCys and pGAD-C1-Rab11-FIPs-C or pGAD-C1-RBD11 (WT/mutants) as described previously (Fukuda et al., 2008; 2011). Yeast cells on the selection medium (SC-AHLW: synthetic complete [SC] medium lacking adenine, histidine, leucine and tryptophan) were incubated at 30°C for around 1 week.

### Cell culture and transfections

COS-7 cells and MDCK cells (parental and *Rab*-KO MDCK-II cells; RIKEN BioResource Center, cat#: RCB5112, RCB5114, RCB5125, RCB5139, and RCB5148) (see Homma et al., 2019; *Rab11*-KO#27) were cultured at 37°C in Dulbecco’s modified Eagle’s medium (DMEM) (FUJIFILM Wako Pure Chemical Corporation) supplemented with 10% fetal bovine serum, 100 units/mL penicillin G, and 100 μg/mL streptomycin in a 5% CO_2_ incubator. One day after plating COS-7 cells in a 6-cm dish (3 × 10^5^ cells), plasmids were transfected into the cells by using Lipofectamine 2000 (Thermo Fisher Scientific) according to the manufacturer’s instructions.

### Retrovirus production and infection of MDCK cells

For retrovirus production, Plat-E cells (a kind gift from Dr. Toshio Kitamura, The University of Tokyo, Tokyo, Japan) (Morita et al., 2000) were plated on a 35mm-dish (4 × 10^5^ cells/dish) and incubated for 24 hr. The cells were transiently transfected with pMRX and pLP/VSVG plasmids (Thermo Fisher Scientific) by Lipofectamine 2000. After 24 hr, the medium was replaced with fresh medium, and the cells were cultured for an additional 24 hr. The medium was then collected and centrifuged at 17,900 × *g* for 3 min to remove debris. The virus-containing medium was added to the MDCK cell culture with 8 μg/mL polybrene. Uninfected cells were removed by treatment with 1 μg/mL puromycin.

### MDCK 3D cyst formation

MDCK cells were suspended in the culture medium containing 12 mM HEPES, pH 7.2 and 2 mg/mL collagen I on ice. The mixture was then dispensed into the 24-well plate and maintained at 37°C for 1 hr. After adding 2 mL of culture medium to each well, the cells were cultured for 7 or 8 days. Then, 2 μg/mL of Dox (Figure 5C and 5D) or 250 nM D/D solubilizer (Figure 5E and 5F) was added to the culture medium for the times indicated in each figure.

### Immunofluorescence analysis

Cells were fixed with 10% trichloroacetic acid (TCA; for MDCK cysts) or 4% paraformaldehyde (PFA; for other cell cultures), permeabilized with 0.1% Triton X-100 in phosphate-buffered saline (PBS) for 3 min (for MDCK cysts) or 50 μg/mL digitonin in PBS for 5 min (for other cell cultures), and incubated with a blocking solution (1% bovine serum albumin in PBS) at room temperature for 1 hr (for MDCK cysts) or 20 min (for other cell cultures). The cells were then incubated for 1 hr at room temperature with primary antibodies, i.e., anti-Rab11 (1/300 dilution), anti-GM130 (1/500 dilution), anti-EEA1 (1/500 dilution), anti-LBPA (1/500 dilution), anti-TfR (1/500 dilution), anti-LAMP2 (1/500 dilution), anti-GFP (1/2000 dilution), and anti-ezrin antibody (1/300 dilution), then for 1 hr at room temperature with Alexa Fluor 555^+^-conjugated anti-rabbit IgG together with DAPI. Only the fixation step was performed on the samples to be examined for EGFP fluorescence alone (Figure 3D). All samples were examined through a confocal fluorescence microscope (Fluoview 1000; Olympus, Tokyo, Japan) equipped with a Plan-Apochromat 100×/1.45 oil-immersion objective lens.

### Immunoblotting

Protein extracts were obtained from cells that had been lysed with a lysis buffer (50 mM HEPES-KOH, pH7.2, 150 mM NaCl, 1% Triton-X100, 1 mM EDTA, and protease inhibitor cocktail [Roche, Basel, Switzerland]) and boiled for 5 min with an SDS sample buffer. Proteins were separated by 16% SDS-polyacrylamide gel electrophoresis (PAGE) and transferred to a polyvinylidene difluoride membrane (Merck Millipore, Burlington, MA) by electroblotting. The blots were blocked for 30 min with 1% skimmed milk in PBS containing 0.1% Tween-20, and after incubation for 1 hr with primary antibodies, they were incubated for 1 hr with appropriate HRP-conjugated secondary antibodies. The entire procedure was performed at room temperature. Chemiluminescence signals were visualized by means of the Immobilon Western Chemiluminescent HRP substrate (EMD Millipore, Burlington, MA) and detected with a chemiluminescence imager (ChemiDoc Touch; Bio-Rad, Hercules, CA).

### Direct binding assays

GST-Rab11A(CA), GST-Rab11A(CN), and control GST alone were expressed in *E. coli* JM109 and purified with gluthathione-Sepharose beads (GE Healthcare) by the standard protocol. For GTP/GDP loading, 10 μg of GST-Rab11A(CA) or GST-Rab11A(CN) was incubated for 20 min at 4°C with 100 μl of 50 mM HEPES-KOH, pH 7.2, 150 mM NaCl, 2.5 mM MgCl_2_, and 0.1% Triton X-100, and then with 1 μl each of 1M MgCl_2_ (final 10 mM) and 50 mM GTPγS (final 0.5 mM) or 100 mM GDP (final 1 mM). COS-7 cells (6-cm dish) transiently expressing T7-tagged Rab11 FIP2-C or RBD11 were lysed for 1 hr at 4°C with 400 μl of 50 mM HEPES-KOH, pH 7.2, 250 mM NaCl, 1 mM MgCl_2_, 1% Triton X-100, and 1×protease inhibitor cocktail (Roche). After centrifugation at 20,000 ×*g* for 10 min, the supernatant was recovered and incubated for 1 hr at 4°C with anti-T7 tag-antibody-conjugated agarose beads (wet volume 30 μl). The beads coupled with T7-tagged proteins were washed three times with 400 μl of 50 mM HEPES-KOH, pH 7.2, 150 mM NaCl, 1 mM MgCl_2_, and 0.1% Triton X-100 (washing buffer). The beads coupled with purified T7-tagged proteins were incubated for 1 hr at 4°C with 100 μl of the solution containing GTPγS-loaded GST-Rab11A(CA) or GDP-loaded GST-Rab11A(CN) described above. After washing the beads with 400 μl of the washing buffer three times, proteins bound to the beads were analyzed by performing 15% SDS-PAGE and staining with Coomassie Brilliant Blue R-250.

### Phylogenetic analysis

The amino acid sequences of the C-terminal region of Rab11-FIPs (102 AA of Rab11-FIP1, 124AA of Rab11-FP2, 100 AA of Rab11-FIP3, 99 AA of Rab11-FIP4, and 101 AA of Rab11-FIP5) were aligned by using the ClustalW software program (version 2.1; available at http://clustalw.ddbj.nig.ac.jp/top-e.html) set at the default parameters and their phylogenetic tree was drawn by the neighbor-joining method.

### Statistical analysis

One-way ANOVA and Tukey’s test or the two-sided Student’s unpaired *t*-test were used to perform the statistical analysis, and *p* <0.05 was used as the criterion for statistical significance.

## ACKNOWLEDGMENTS

We thank Drs. Toshio Kitamura and Shoji Yamaoka for kindly donating materials, Kazuyasu Shoji for technical assistance, and members of the Fukuda laboratory for valuable discussions.

## AUTHORS CONTRIBUTIONS

Conceptualization, F.O., T.M., and M.F.; Methodology, S.H., and Y.H.; Investigation, F.O., T.M., and M.F.; Writing – Original Draft, F.O. and M.F.; Writing – Review & Editing, F.O., T.M., S.H., Y.H., and M.F.; Funding Acquisition, T.M., Y.H., and M.F.; Supervision, T.M. and M.F.

