## Supplementary material for "RBD11, a bioengineered Rab11-binding module for visualizing and analyzing endogenous Rab11": Suppl Figs 1 & 2

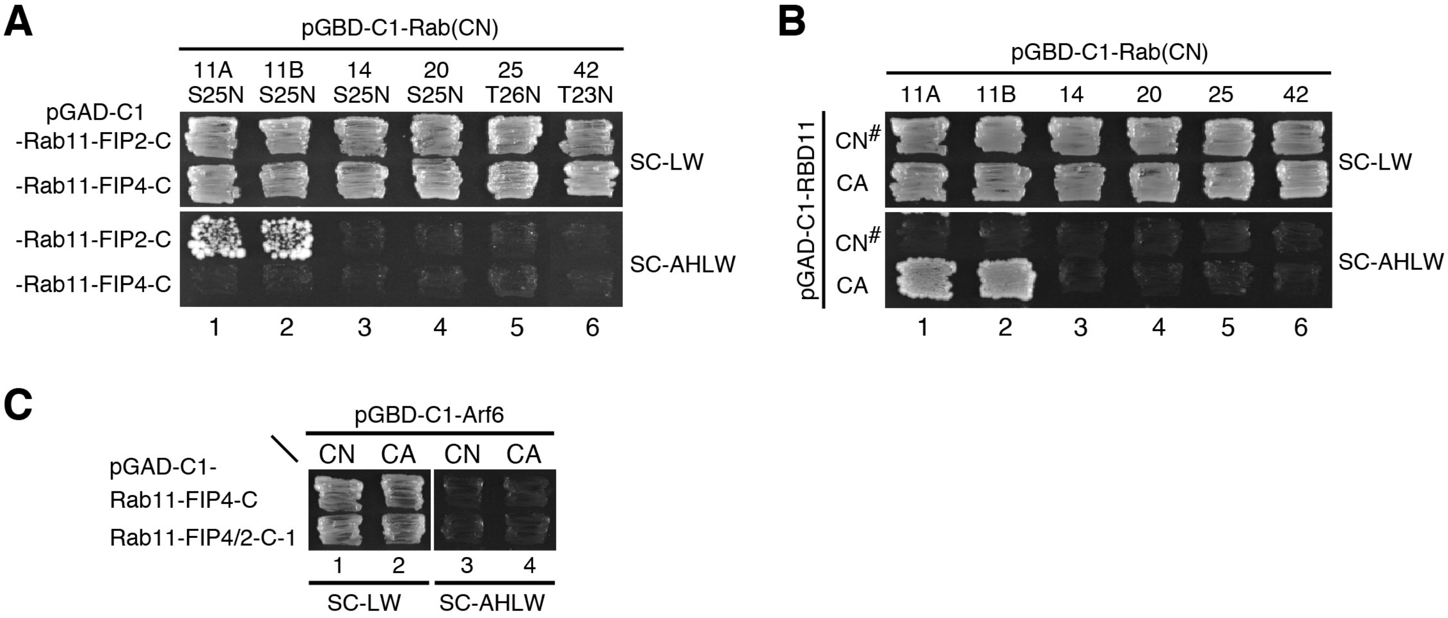


**Figure S1. Rab binding specificity of Rab11-FIP2-C, Rab11-FIP4-C, and RBD11, and Arf6 binding activity of RBD11 (related to Figures 1 and 3)**

(A) Rab binding specificity of Rab11-FIP2-C and Rab11-FIP4-C as determined by yeast two-hybrid assays. Rab11-FIP2/4-C were subcloned into the pGAD-C1 vector, and it was then transformed into yeast cells expressing pGBD-C1 vector carrying a constitutively negative form, which contains a Ser/Thr-to-Asn substitution, of each Rab. Yeast cells were grown on SC-LW (growth medium) and SC-AHLW (selection medium).

(B) Rab binding specificity of RBD11 as determined by yeast two-hybrid assays. CN# indicates the constitutively negative mutants with a Ser/Thr-to-Asn substitution shown in (A).

(C) Arf6 binding activity of Rab11-FIP4-C and Rab11-FIP4/2-C1 as determined by yeast two-hybrid assays. The pGAD-C1 vector carrying pGAD-C1-Rab11-FIP4-C (or -Rab11-FIP4/2-C1) was transformed into yeast cells expressing pGBD-C1 vector carrying a CN or CA form of Arf6.

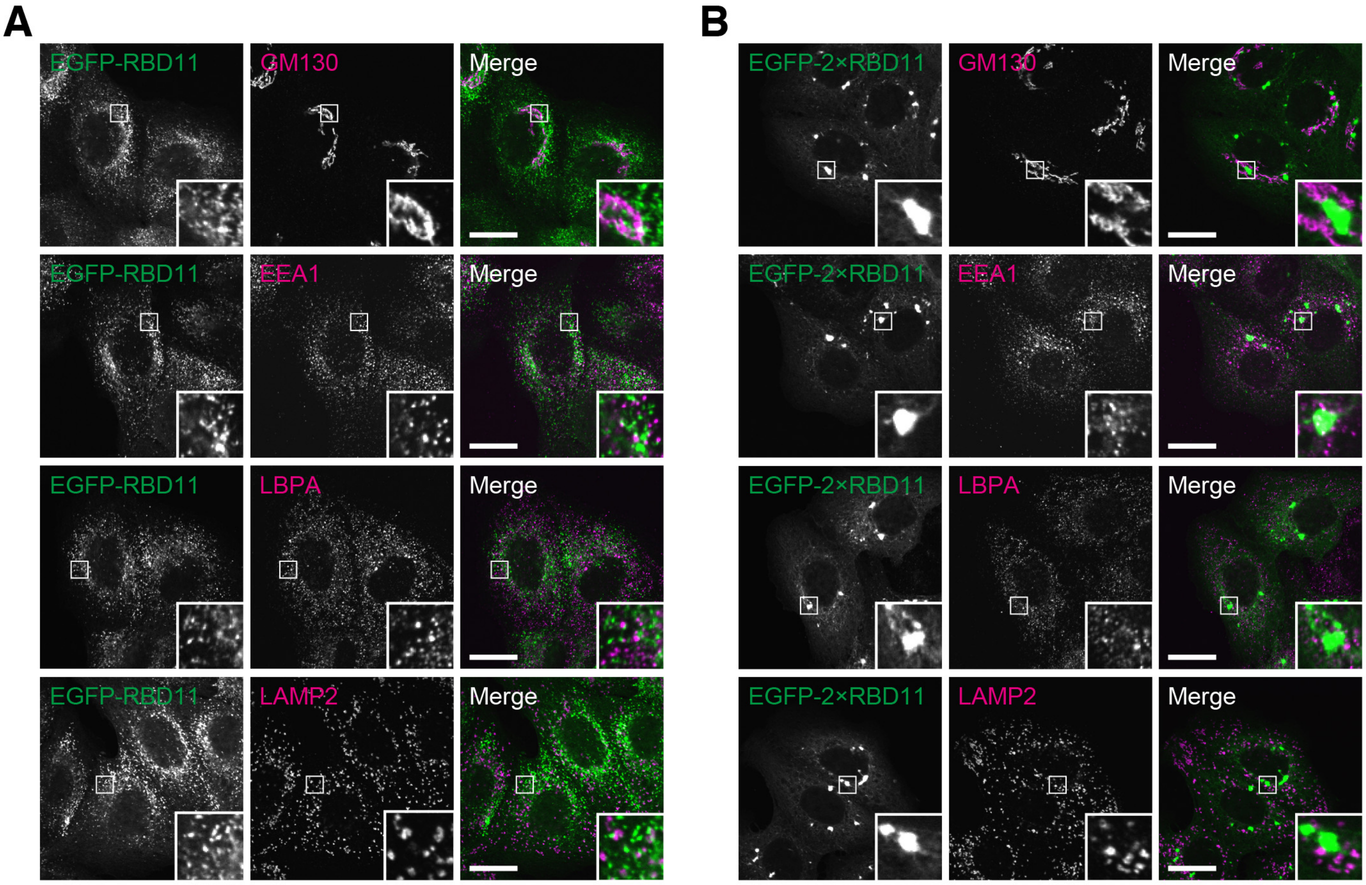


**Figure S2. Neither RBD11 nor 2×RBD11 colocalized with the Golgi, early endosomes (EEs), late endosomes (LEs), or lysosomes (LYs) (related to Figures 3 and 4)**

(A) MDCK cells stably expressing EGFP-RBD11 (green) were fixed with 4% PFA and immunostained with antibodies against GM130 (Golgi marker), EEA1 (EE marker), LBPA (LE marker), and LAMP2 (LY marker) (red). Scale bars, 20 μm.

(B) MDCK cells stably expressing EGFP-2×RBD11 (green) were fixed with 4% PFA and immunostained with the antibodies shown in (A) (red). Scale bars, 20 μm.
